# Correcting statistical bias in correlation-based kinship estimators

**DOI:** 10.1101/2021.01.13.426515

**Authors:** Wei Jiang, Xiangyu Zhang, Siting Li, Shuang Song, Hongyu Zhao

**Affiliations:** Department of Biostatistics, School of Public Health, Yale University, New Haven, Connecticut, United States of America; Department of Statistics, University of Science and Technology of China, Hefei, Anhui, China; Department of Biomedical Data Science, Geisel School of Medicine, Dartmouth College, Hanover, New Hampshire, United States of America; Center for Statistical Science, Tsinghua University, Beijing, China; Department of Industrial Engineering, Tsinghua University, Beijing, China

## Abstract

Accurate estimate of relatedness is important for genetic data analyses, such as association mapping and heritability estimation based on data collected from genome-wide association studies. Inaccurate relatedness estimates may lead to spurious associations and biased heritability estimations. Individual-level genotype data are often used to estimate kinship coefficient between individuals. The commonly used sample correlation-based genomic relationship matrix (scGRM) method estimates kinship coefficient by calculating the average sample correlation coefficient among all single nucleotide polymorphisms (SNPs), where the observed allele frequencies are used to calculate both the expectations and variances of genotypes. Although this method is widely used, a substantial proportion of estimated kinship coefficients are negative, which are difficult to interpret. In this paper, through mathematical derivation, we show that there indeed exists bias in the estimated kinship coefficient using the scGRM method when the observed allele frequencies are regarded as true frequencies. This leads to negative bias for the average estimate of kinship among all individuals, which explains the estimated negative kinship coefficients. Based on this observation, we propose an unbiased estimation method, UKin, which can reduce the bias. We justify our improved method with rigorous mathematical proof. We have conducted simulations as well as two real data analyses to demonstrate that both bias and root mean square error in kinship coefficient estimation can be reduced by using UKin. Further simulations indicate that the power in association mapping can also be improved by using our unbiased kinship estimates to adjust for cryptic relatedness.

**Author summary:** Inference of relatedness plays an important role in genetic data analysis. Many methods have been proposed to estimate kinship coefficients, including the commonly used genomic relationship matrix method. However, a substantial proportion of the kinship coefficients estimated by this method are negative, which is difficult to interpret. In this paper, through mathematical derivation, we show that there indeed exists a negative bias in this approach. To correct for this bias, we propose a new kinship coefficient estimation method, UKin, which is unbiased without requiring extra genetic information nor added computational complexity. The better performance of UKin in reducing bias and root mean squared error is demonstrated through theory, simulations and analysis of data from the young-onset breast cancer and familial intracranial aneurysm studies.

## Introduction

Accurate estimation of relatedness among individuals is important in genetic data analysis. For example, in both population-based and family-based genome-wide association studies (GWAS) with uncertain relationships among study subjects, it is critical to appropriately account for cryptic relatedness because incorrect estimates can decrease power and inflate false positive rates of association tests [1–3]. Several methods have been proposed to adjust for relatedness in GWAS, such as introducing a genomic relationship matrix (GRM) as an augment into well-developed linear mixed model (LMM) [4–7]. It has been demonstrated that proper consideration of genetic relatedness can also benefit heritability estimation based on GWAS data in the presence of pedigree structures [8, 9].

In order to adjust for cryptic relatedness in genetic studies like association mapping and heritability estimation, individual-level genotype data are often used to estimate pairwise kinship coefficients. The sample correlation-based genomic relationship matrix (scGRM) method estimates kinship coefficient by calculating the average sample correlation coefficient among all genetic variants, in which the observed allele frequencies are used for the calculation of both expectation and variance of genotypes [10–12]. We note that most association mapping and heritability estimation packages use this method as their default setting for calculating GRM, such as GCTA, GEMMA and FaSTLMM [6, 8, 13]. Although this method is widely used, researchers have noted that a substantial proportion of the estimated kinship coefficients are negative. As kinship coefficient is defined to be a positive number (see in Materials and Methods), it is difficult to interpret these negative estimates [14–16].

In this paper, through mathematical derivation, we first show that there indeed exists bias in the estimated kinship coefficients using the scGRM method. The bias exists because the observed allele frequencies are regarded as true frequencies. We also prove analytically that the bias essentially results in a negative average for all estimates, which explains the large proportion of negative values. Based on this observation, we propose an improved kinship estimation method, UKin, which can remove bias. We provide a mathematical proof for the unbiasedness of the UKin estimator. Simulations and real data analyses also demonstrate that both bias and root mean square error (RMSE) can be reduced by replacing the scGRM method with our UKin method. For real data analyses, we apply our method to two studies, young-onset breast cancer (BC) and familial intracranial aneurysm (FIA), which have pedigree information to evaluate our results. Finally, as an application of our method in association mapping, we conduct a simulation study to show the power of detecting genetic associations can be improved by correcting cryptic relatedness using our unbiased kinship estimates.

The paper is organized as follows. In the Results section, we evaluate the performance of UKin through two simulations and two real data sets in BC and FIA to validate our theoretical derivation and demonstrate the effectiveness of UKin estimator in reducing bias and RMSE. In the Materials and Methods section, we present the theoretical details which show the scGRM method is biased, propose our UKin estimation method and give the correctness proof, as well as its connection with the scGRM estimator. Lastly, we conclude with a simulation study in the Discussion section to demonstrate that UKin method can further improve the power of association mapping. Technical details such as mathematical derivations are provided in S1 Appendix.

## Results

### 1. Simulation experiments

#### An illustrative example

We start our discussion with a simple but extreme example. In this experiment, we assumed that there were 500 full siblings from the same family. Although unlikely to exist in reality, this example serves as a good illustration of our theoretical derivation. As every two individuals selected from the same family were full siblings, the true value of their kinship coefficient should be 0.25 (see in Table 5). However, following Property 3 in the Materials and Methods section, their average kinship coefficient estimated by scGRM, denoted by 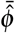, should have the expectation:

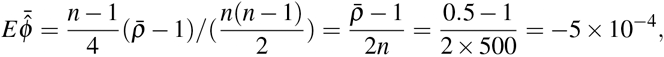

where *n* is the sample size and 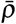 is the average of their true genetic correlation coefficients. Property 1 together with Table 5 in the Materials and Methods section suggest that 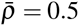 for full siblings.

This result shows the unexpected phenomenon that although all individuals in our simulated samples are full siblings to each other, the average of the estimated kinship coefficients has a negative value. To illustrate Property 3 in practice, we simulated 200 unrelated families each consisting of 500 full siblings with the method provided by the package CorBin [17]. Each individual was genotyped at 10,000 single nucleotide polymorphisms (SNPs). Following the scGRM method and the UKin method proposed in Materials and Methods, we estimated pairwise kinship coefficients and calculated their mean values, respectively. The histograms of these estimated average kinship coefficients are shown in Fig 1. From this plot, we could see the distribution of average kinship estimated by the scGRM method centered around *−*5 *×* 10^*−*4^, which is consistent with our expectation from the analytical results. By contrast, the UKin approach performed better in dealing with this extreme situation, with the average estimates centered at 0.25, the true value of pairwise kinship coefficient for full-sibling pairs. Besides, from Fig 1 we could observe that the two distributions have similar shapes, which could be explained by Equation 5 in Materials and Methods which suggests that unbiased estimator of correlation coefficient 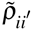 could be expressed as a linear combination of the scGRM estimators 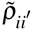. Considering there were 500 full siblings from the same family, we calculated the average on both sides of Equation 5 among all the simulated individual pairs, which is 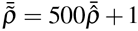, where 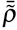 and 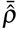 represent the average of correlation coefficients between full siblings from the same family, estimated by the UKin method and the scGRM method respectively, i.e.

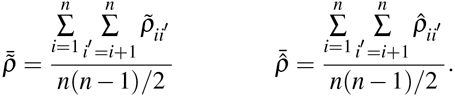

**Fig 1.**
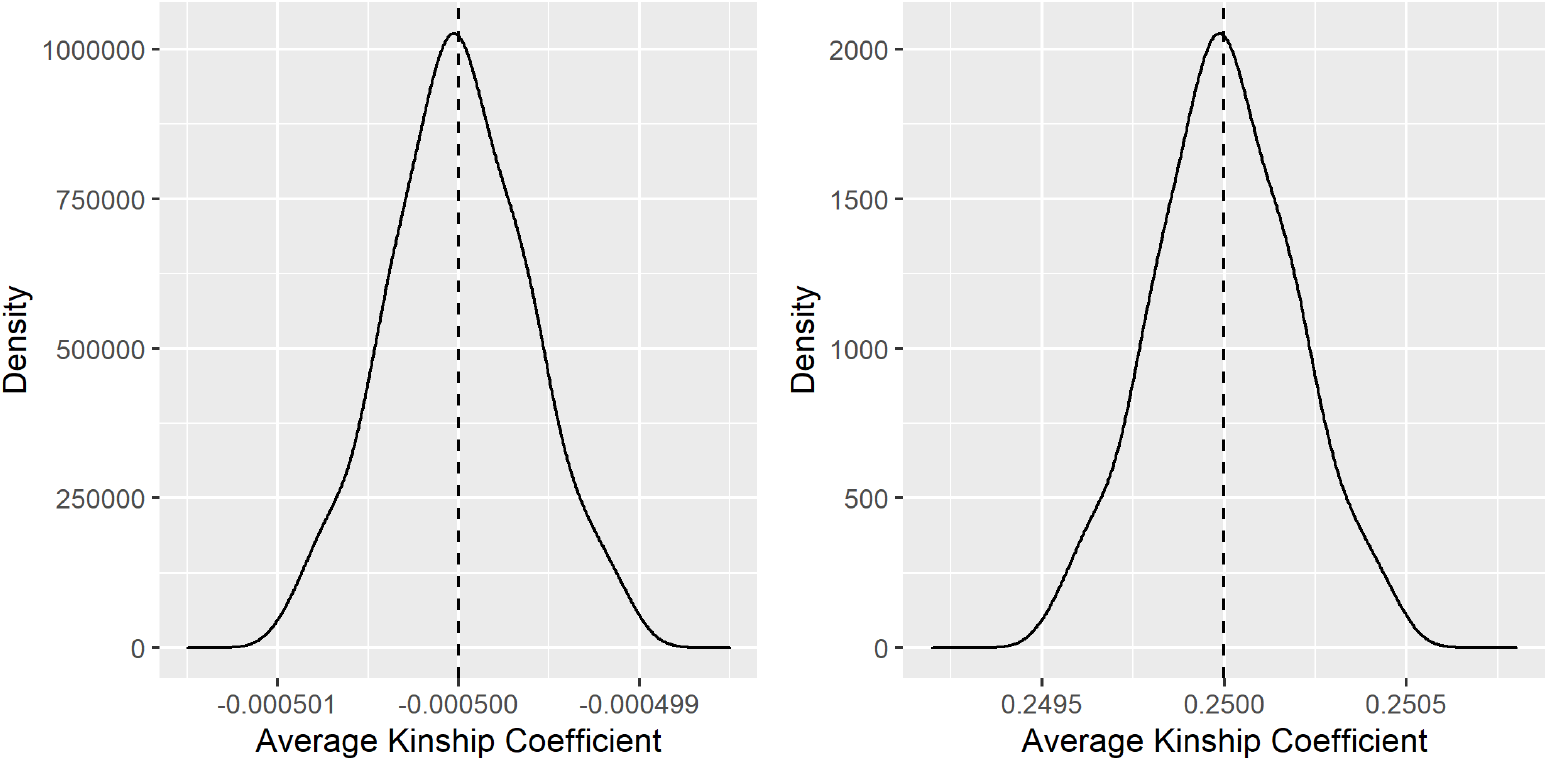
Distribution of average kinship coefficients estimated by the scGRM (left) and UKin (right) methods in this extreme example. Two hundred unrelated families each consisting of 500 full siblings were simulated, with each sibling genotyped at 10,000 SNPs. The averages of kinship coefficients among all individual pairs from the same family were calculated and the distribution of these averages is displayed. The true value of kinship coefficient between full siblings is 0.25. The vertical dashed line in each plot corresponds to the mean value of these averages estimated by the corresponding method.

As there was a linear relationship between kinship coefficient and correlation coefficient (see Property 1 in Materials and Methods), the distributions of the average kinship coefficients estimated by the two methods should have the same shape.

#### A more general simulation

To evaluate the performance of the UKin method in kinship coefficient estimation and to compare it with the scGRM estimate in a more general situation, we performed the following simulations in which population homogeneity was assumed. To include different kinds of relationships in our experiment, we simulated 6,000 people including 1,000 pairs with kinship coefficient 0.125, 1,000 pairs with coefficient 0.25, and 500 pairs with coefficient 0.5. For simplicity, different relative pairs were set to be unrelated. In addition, we also included 1,000 people who had no relationship with other individuals. For each subject, genotype data were generated for 10,000 random and independent SNPs. The minor allele frequencies (MAFs) of genotyped variants were drawn uniformly from [0.05, 0.5].

With the UKin and scGRM estimators, we estimated kinship coefficients between all simulated individual pairs and divided those coefficients into four groups according to their true relationships. Fig 2 shows the distribution of the estimated kinship coefficients in each group respectively. As shown in this plot and summarized in Table 1, for groups with true kinship coefficient 0.25 and 0.5, our UKin method achieved lower RMSE than the scGRM method in estimating kinship coefficients, while the opposite was true for the independent pairs. For the group consisting of pairs having kinship coefficients 0.125, the two methods had similar performance.

**Table 1.** Comparison of UKin and scGRM in biases and RMSEs

| True Value | Bias from True Value($\times 10^{-5}$) | | Root Mean Square Error($\times 10^{-3}$) | |
| --- | --- | --- | --- | --- |
|  | UKin | scGRM | UKin | scGRM |
| 0.000 | **3.943** | -11.81 | 6.915 | **5.000** |
| 0.125 | **1.543** | -7.486 | 5.724 | **5.668** |
| 0.250 | **3.414** | -13.80 | **4.537** | 6.329 |
| 0.500 | **0.000** | -22.82 | **0.000** | 7.431 |

**Fig 2.**
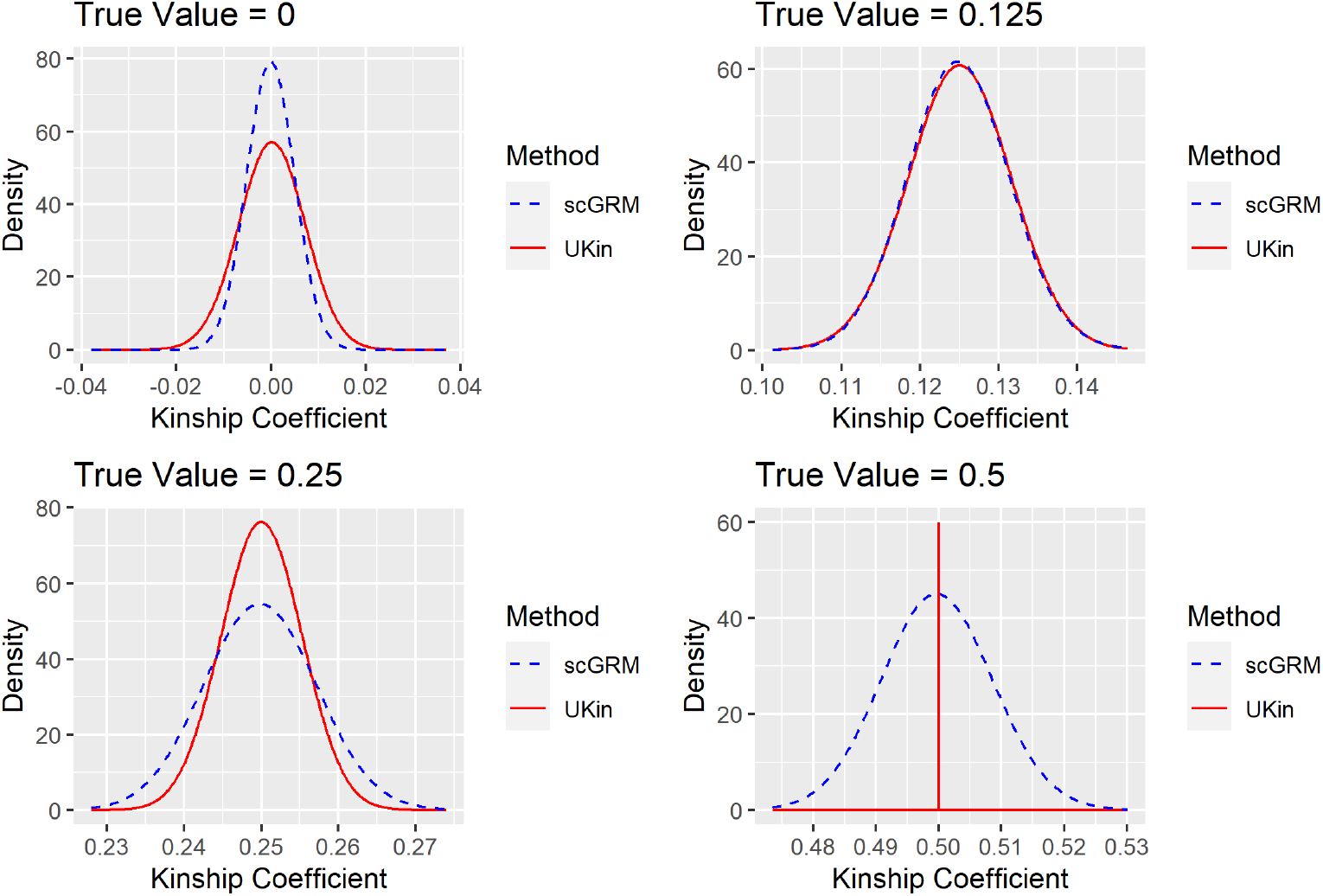
Distributions of kinship coefficients estimated by the UKin method and the scGRM method in our simulation study including 6,000 individuals with different relationships. The four plots correspond to the four groups divided by the true value of estimated kinship coefficients.

Although Fig 2 clearly demonstrates the RMSE for the two methods, it is difficult to compare their biases from the plots. More detailed comparisons are shown in Table 1. As shown in the second column of this table, UKin always performed better than scGRM when we compared the mean values of estimated kinship coefficients, as the results of the UKin method were closer to true values for all four groups. Besides, results in the third column of Table 1 show that UKin could reduce RMSE for close relatives, which is consistent with the conclusion we get from Fig 2. Furthermore, UKin shows a downward trend of RMSE with increasing true kinship coefficients, while scGRM is completely on the opposite. It is also notable that when we consider individual pairs with kinship coefficient 0.5, i.e. Monozygotic twins (M-Z twins), both bias and RMSE are extremely close to zero if we utilize UKin to estimate.

### 2. Real data analyses

#### The Young-Onset Breast Cancer Study

To demonstrate our unbiased method could get more accurate results in estimating kinship coefficients, we applied the UKin method to real data from a family-based study of genes and environment in young-onset BC (*dbGaP Study Accession: phs000678.v1.p1*). This study recruited families from the US and Puerto Rico with a daughter who was recently diagnosed with breast cancer and another unaffected daughter. For each family, only the diseased daughter and her unaffected full sister were genotyped for analysis. As for data quality control, we removed individuals with more than 10% missing genotypes as well as SNPs with a missing genotype rate greater than 5% or a minor allele frequency less than 5%. After further removing individuals with missing phenotypes, we got 1,983 subjects (1,458 cases and 525 controls) with 614,310 variants in total. The processed data included 511 pairs of full sisters, with one affected by breast cancer. We assumed individuals from different families were unrelated, then the true values of all estimated kinship coefficients should be either 0.25 (511 full sister pairs) or 0 (all the other individual pairs).

We first applied the scGRM method to estimate the kinship coefficients, which had poor performance. For the 511 within-family pairs (full sister pairs), only 473 pairs were estimated to have kinship coefficients between 2^*−*5*/*2^ and 2^*−*3*/*2^, which means 7.4% of full sisters were incorrectly inferred to be other kinds of relative pairs. Estimation of 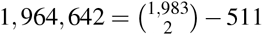 between-family pairs with the scGRM method also performed poorly, as 0.45% unrelated pairs between families were misspecified as 1st-degree relative pairs (such as sibling pairs), 2nd-degree relative pairs (such as half-sibs, avuncular pairs and grandparent-grandchild pairs) or 3rd-degree relative pairs (such as first cousins). In contrast, our UKin method had more accurate estimates with 501 out of 511 (98.0%) full sisters pairs correctly estimated and only 177 unrelated pairs (less than 0.01%) were misspecified as 3rd-degree relative pairs (Table 2 and Table 3).

**Table 2.**
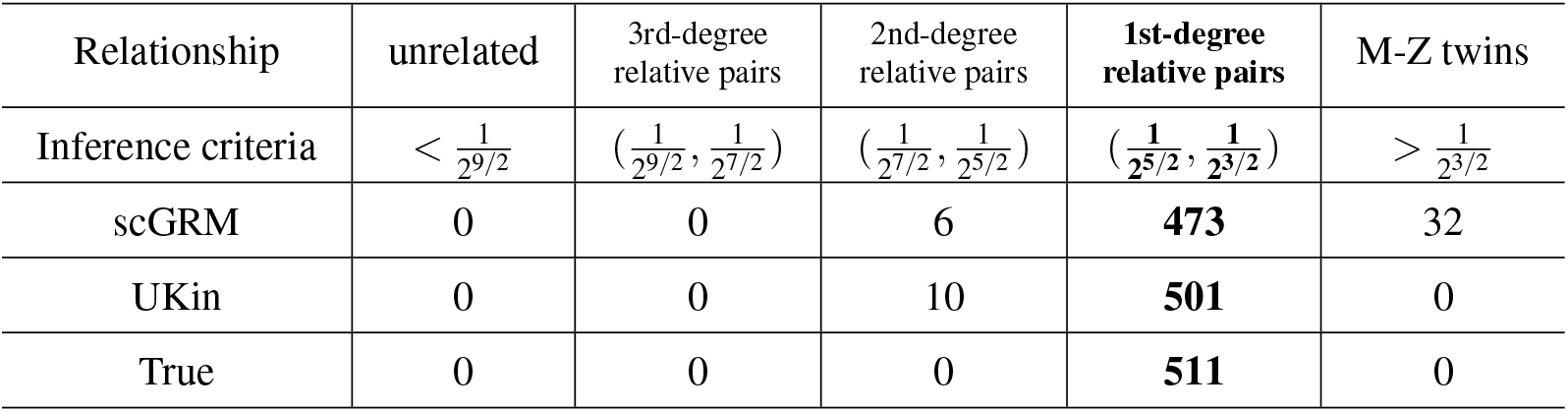
Distribution of estimated kinship coefficients of 511 full siblings in the two sister data studying young-onset breast cancer

| Relationship | unrelated | 3rd-degree relative pairs | 2nd-degree relative pairs | 1st-degree relative pairs | M-Z twins |
| --- | --- | --- | --- | --- | --- |
| Inference criteria | $< \frac{1}{2^{9/2}}$ | $(\frac{1}{2^{9/2}}, \frac{1}{2^{7/2}})$ | $(\frac{1}{2^{7/2}}, \frac{1}{2^{5/2}})$ | $(\frac{1}{2^{5/2}}, \frac{1}{2^{3/2}})$ | $> \frac{1}{2^{3/2}}$ |
| scGRM | 0 | 0 | 6 | **473** | 32 |
| UKin | 0 | 0 | 10 | **501** | 0 |
| True | 0 | 0 | 0 | **511** | 0 |

**Table 3.**
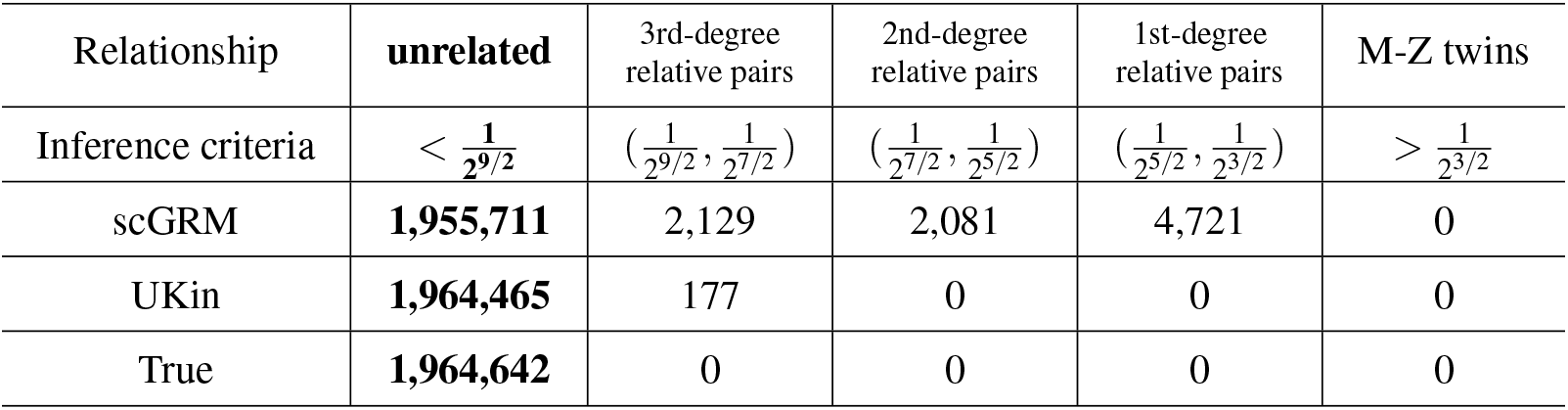
Distribution of estimated kinship coefficients of 1,964,642 unrelated individual pairs in the two sister data studying young-onset breast cancer

| Relationship | unrelated | 3rd-degree relative pairs | 2nd-degree relative pairs | 1st-degree relative pairs | M-Z twins |
| --- | --- | --- | --- | --- | --- |
| Inference criteria | $< \frac{1}{2^{9/2}}$ | $(\frac{1}{2^{9/2}}, \frac{1}{2^{7/2}})$ | $(\frac{1}{2^{7/2}}, \frac{1}{2^{5/2}})$ | $(\frac{1}{2^{5/2}}, \frac{1}{2^{3/2}})$ | $> \frac{1}{2^{3/2}}$ |
| scGRM | **1,955,711** | 2,129 | 2,081 | 4,721 | 0 |
| UKin | **1,964,465** | 177 | 0 | 0 | 0 |
| True | **1,964,642** | 0 | 0 | 0 | 0 |

The histograms of kinship coefficients estimated by scGRM and UKin for all pairs (including both full sister pairs and unrelated pairs) in the BC study are given in Fig 3. To make the comparison more clearly, we only took individual pairs with estimated kinship coefficients between 2^*−*5^ and 2^*−*1.5^ into consideration. It is obvious that the histogram corresponding to scGRM contains more pairs with estimated kinship coefficients larger than 2^*−*7*/*2^. From previous analysis, we know most of them are misspecified unrelated pairs. In contrast, our approach is much less likely to make such mistakes. Besides, the UKin histogram shows a peak centered close to 0.25 and has a distinct separation from estimated kinship coefficients near zero. However, scGRM does not work well in this aspect because the distribution of non-zero kinship coefficients is centered around 0.23 and has an obvious distribution overlap with unrelated pairs, which suggests that UKin performs better in separating relatives from unrelated pairs. Furthermore, if we only consider the 511 full sister pairs, the true kinship coefficient should be However, the average and mean square error (MSE) of the kinship coefficients estimated by scGRM were 0.263 and 4.03 *×* 10*−*3, respectively. In contrast, the corresponding UKin results were 0.248 and 6.72 *×* 10*−*4, respectively. To visualize the difference between UKin and scGRM, we also draw the scatter plot of the estimated kinship coefficients for the 511 full sister pairs between the two methods (Fig 4). The scatter plot demonstrates that while the distribution of UKin estimates is more concentrated at its true value, scGRM tends to overestimate the kinship coefficients for many full sister pairs. These results clearly show the better performance of UKin than scGRM.

**Fig 3.**
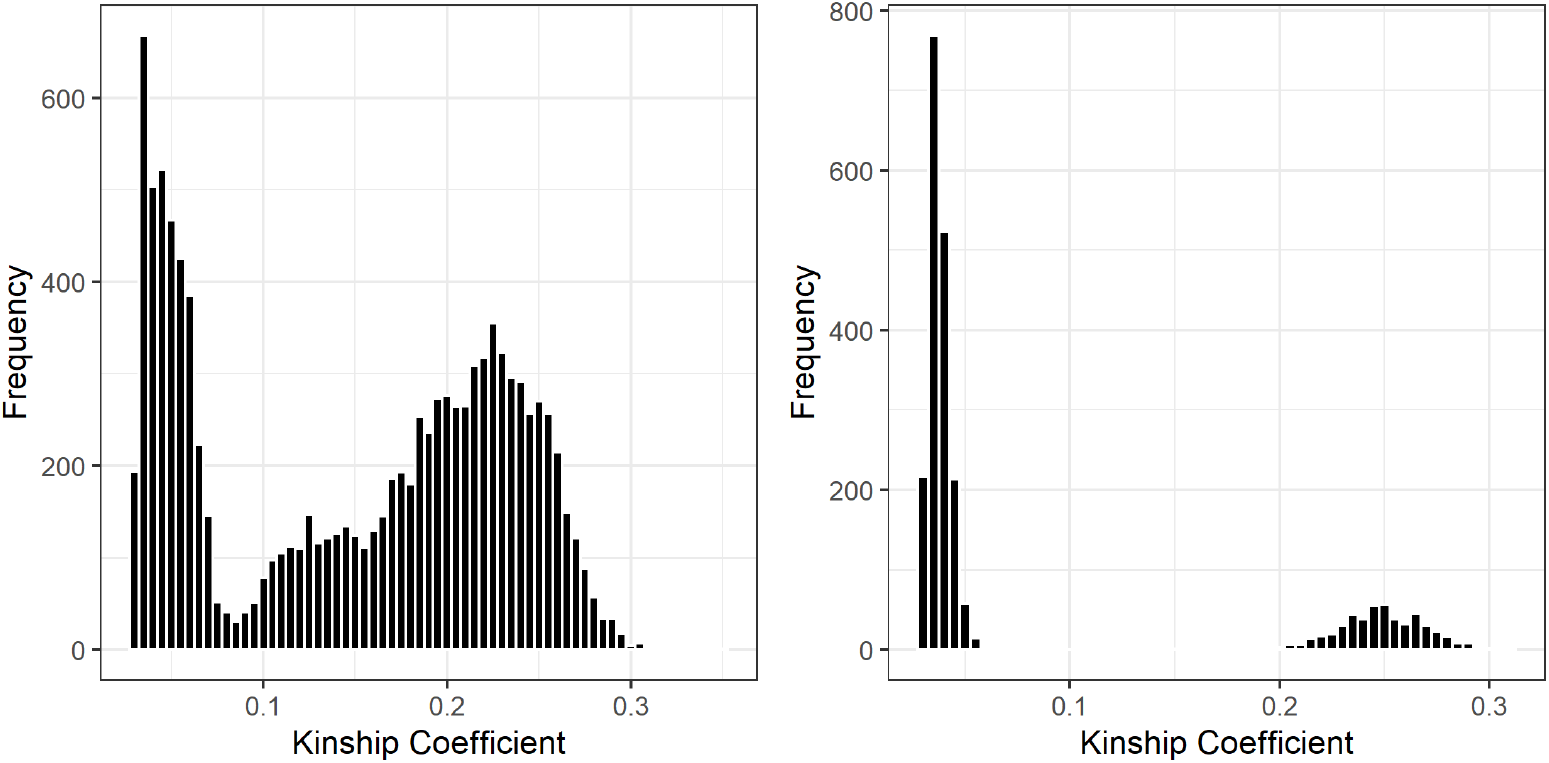
Comparison of distributions of kinship coefficients estimated by scGRM (left) and UKin (right) in breast cancer study. This study genotyped 1,983 individuals at 614,310 variants. Pairwise relationships in this dataset included 511 full sister pairs from irrelevant families and unrelated pairs. In this figure, we only considered estimated kinship coefficients between 2^*−*5^ and 2^*−*1.5^. Class interval of the histogram for each method is set to be 0.005.

**Fig 4.**
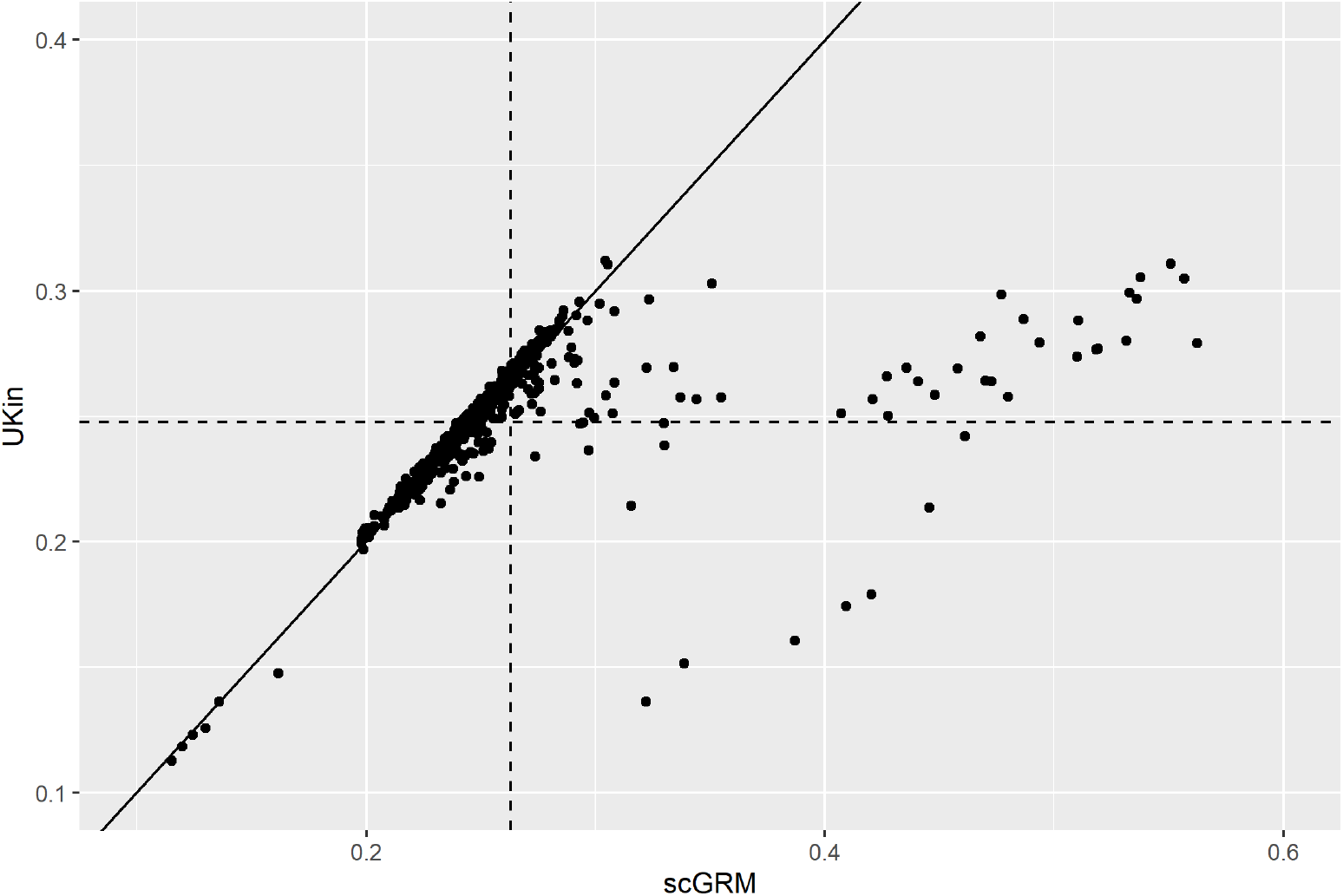
The scatter plot of the estimated kinship coefficients between the UKin and scGRM methods. For this plot we only consider the 511 full sister pairs in the BC data set. The oblique solid line stands for the equation *y* = *x*, while the vertical and horizontal dashed lines correspond to the mean values of scGRM and UKin estimates, respectively.

#### The Familial Intracranial Aneurysm Linkage Study

To further investigate the effectiveness of the UKin method in kinship coefficient estimation, we applied UKin to infer pedigree structure using genotype data from the FIA linkage study (*dbGaP Study Accession: phs000293.v1.p1*). This study recruited 400 families with multiple individuals who have an intracranial aneurysm (IA) through 23 (25) referral centers throughout North America, Australia, and New Zealand that represent 35 (40) recruitment sites. After a standardized procedure of quality control and discarding subjects with missing phenotype, we obtained 990 individuals from 371 families and each of them was genotyped at 5,505 SNPs. In this FIA dataset, the confirmed relationships include 137 1st-degree relative pairs (including 19 full siblings and 118 parent-child pairs).

We compared the performance of UKin and scGRM in identifying these 1st-degree relative pairs and estimating their kinship coefficients. UKin was able to correctly recognize all the 137 1st-degree pairs (with estimated kinship coefficients between 2^*−*2.5^ and 2^*−*1.5^), while scGRM misspecified one parent-child pair as monozygotic twins, with an estimated kinship coefficient of 0.442. The histograms of the kinship coefficients of these 137 individual pairs estimated by the two methods (Fig 5) indicate that unbiased estimations are more concentrated, taking values between 0.21 and 0.3. However, the distribution of scGRM estimations is more dispersed, including a distinct outlier. This fact is more clearly shown in the scatter plot including all the 137 1st-degree pairs in the IA data set (Fig 6). We further calculated the bias from the true value (0.25) and RMSE of the estimated coefficients for each estimator. As summarized in Table 4, the estimation bias of UKin was 1/6 of the bias estimated by scGRM, while the RMSE of UKin was half of scGRM. We also note that scGRM misspecified 15 parent-child pairs or unrelated pairs as MZ twins, while UKin only made five such mistakes which were all included in the misspecified pairs of scGRM. These results demonstrate that our UKin method achieves more accurate outcomes in relationship inference and kinship estimation, even when the number of genotyped SNPs is small.

**Table 4.** Bias and RMSE of estimated kinship coefficients for the 137 pairs of 1st-degree relatives in the FIA study

| Estimation Method | Bias($\times 10^{-3}$) | RMSE($\times 10^{-2}$) |
| --- | --- | --- |
| UKin | 0.667 | 1.175 |
| scGRM | 4.045 | 2.244 |

**Fig 5.**
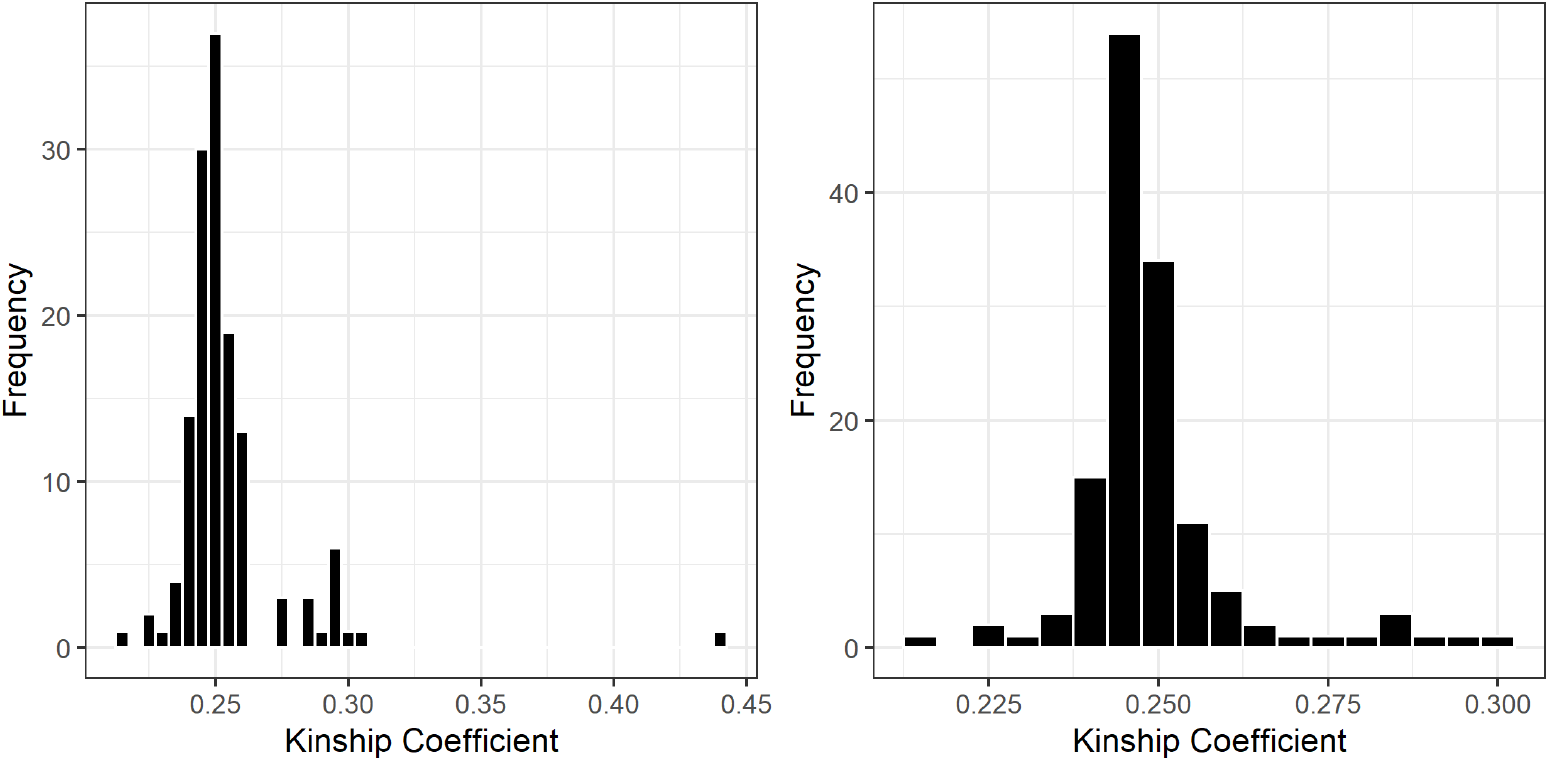
Distributions of estimated kinship coefficients of 1st-degree relatives in the FIA study with scGRM (left) and UKin (right). Among all the 137 1st-degree relative pairs in this dataset, there are 19 full siblings and 118 parent-child pairs. Class interval of the histogram for each method is set to be 0.005.

**Fig 6.**
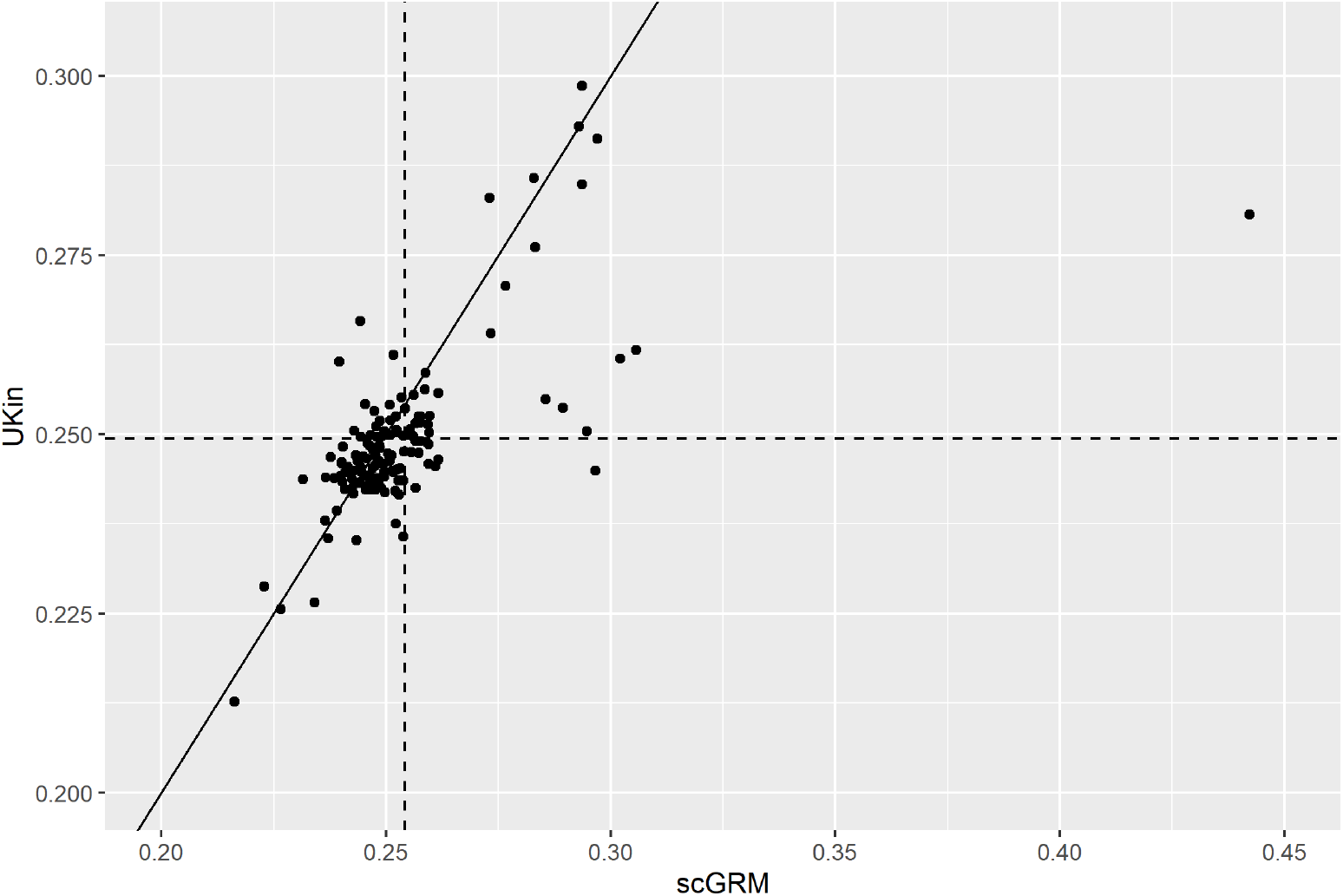
The scatter plot of the estimated kinship coefficients between the UKin and scGRM methods in the FIA study. Only the 137 1st-degree pairs are shown. The oblique solid line stands for the equation *y* = *x*, while the vertical and horizontal dashed lines correspond to the mean values of scGRM and UKin estimates, respectively.

## Discussion

Among the many kinship coefficient estimation methods, the most commonly applied estimator uses dense SNP genotypes and allele frequencies in the samples to calculate average pairwise correlation coefficients among SNPs. Although this method is intuitive and easy to calculate, we prove that it is actually biased because it treats the observed allele frequencies as true frequencies. Through rigorous derivation, we showed that pairwise kinship coefficients estimated by scGRM add up to be a negative value, which explains the phenomenon that a substantial proportion of kinship coefficient estimates are negative.

When conducting large scale estimates of kinship coefficients, the existing bias in scGRM can lead to incorrect inference of relationships, and this problem can be extremely severe if the subjects in the dataset are highly related. Our method, UKin, solved this issue by incorporating genetic information from the whole population to adjust for the bias in the estimated kinship coefficient between every single pair. This unbiased estimator can be expressed as a polynomial of scGRM estimators, and leveraging only information of dense genotypes from the population. As demonstrated by our simulations and applications to the BC and FIA family data, UKin performed better in reducing both estimation bias and RMSEs. For the two sister study, the results suggest that while scGRM could lead to severe spurious inference of relative pairs, UKin rarely made such mistakes. Even when the number of genotyped SNPs was limited for the FIA study, UKin could reduce statistical bias and RMSE while avoiding spurious relationship inference.

In our theoretical derivations and simulation studies, we made assumptions like linkage equilibrium (LE) and absence of inbreeding, that is, genotypes at different markers are independent. During our derivation, we used the same weights for all SNPs, and our simulated datasets were also generated under this assumption. Although there is linkage disequilibrium (LD) in reality, empirical results from the analyses of the BC and FIA family data show that the bias and RMSEs can also be reduced greatly with the application of UKin to real data. To consider the problem of LD in practice, we can give different weights based on LD to these SNPs. Following the approach of Wang (2017) [15], these LD weights **w** = (*w*_1_*, w*_2_*, …, w*_*m*_)^*T*^ can be calculated by solving the following minimization problem:

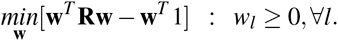

where 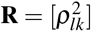 is the matrix of squared LD correlations. Theoretically, this result can be directly applied to UKin by assigning the correlation coefficient at each SNP marker its corresponding weight, which might make our approach adapt to LD situation.

Another assumption throughout our study is a homogeneous population so that the allele frequencies can be calculated once and applied to all subjects. Some methods have been proposed to estimate kinship coefficients in admixed populations, where the assumption of population homogeneity is untenable [11, 18, 19]. However, as most of these methods are based on the scGRM method, they are also likely to be biased estimators, too. How to extend our UKin method to deal with admixed populations is a topic for future studies.

More accurate kinship estimation will improve the performance of different genetic analyses such as association mapping. In recent years, GWAS have seen great success in identifying genetic loci contributing to complex human traits [20, 21]. By studying a genome-wide data set of genetic variants in different individuals, GWAS looks for SNPs correlated with traits in the samples. Accurate specification of familial relationships is expected to bring more powerful association results in GWAS with unknown (or unrecognized) family structure. We have investigated whether association mapping can be improved by applying UKin to account for cryptic relatedness.

We conducted a simulation study to compare the performance of UKin with scGRM in GWAS. In our experiments, we simulated 4,000 samples including 2,000 cases and 2,000 controls. We included subjects with various pairwise kinship coefficients in both cases and controls. More specifically, we simulated 250 1st-degree relative pairs, 250 2nd-degree relative pairs, 250 3rd-degree relative pairs, and 500 unrelated subjects. The total number of SNPs genotyped for each individual was set to be 10,000 and the MAFs of non-risk SNPs were drawn uniformly from [0.05, 0.5]. The proportion of risk SNPs was set at 0.05 or 0.1. For these risk SNPs, a variable following the Gaussian distribution 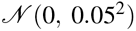 was added to the previous uniform distribution to obtain their MAFs in cases. We set those MAFs below 0.05 or greater than 0.95 to be 0.05 and 0.95, respectively.

We applied GEMMA [13], which was developed to implement the genome-wide mixed model association algorithm for a standard linear mixed model for association analysis. In our simulations, we performed likelihood ratio tests in a univariate LMM for marker association mappings with a single phenotype. PLINK binary file format was [22] adopted as input files containing phenotypes and genetic information. A standardized relatedness matrix file estimated by either scGRM or UKin was included to appropriately account for relatedness among subjects.

We applied GEMMA to analyze the simulated GWAS dataset and selected all SNPs with P-value below the threshold 5 *×* 10^*−*4^. Statistical power and type I error rate were calculated to evaluate the performance of marker association tests when the relatedness matrix used in LMMs was estimated by scGRM and UKin, respectively. The results suggest that the type I error rate was appropriately controlled at a low level (less that 5 *×* 10^*−*4^) for both methods. We compared the power of association mapping which suggests that for the two risk SNP proportions considered (i.e. 0.05 and 0.1), the power of identifying risk variants was always improved after we replaced scGRM with UKin in estimating pairwise kinship coefficients. For example, when the proportion of risk SNPs was set at 0.05, the power was improved from 0.154 to 0.167 by adopting the UKin method. This simulation demonstrates that the application of UKin can improve statistical power while controlling the type I error rate in GWAS. However, further simulations and real data experiments are required to evaluate the advantages of UKin over the scGRM comprehensively, which is the subject of future research.

## Materials and Methods

Alleles are said to be identical by descent (IBD) if they are inherited from a common ancestor. To describe the average amount of IBD sharing at the genome level, we often adopt the concept of kinship coefficient [12]. For two individuals indexed by *a* and *b*, their kinship coefficient, *ϕ*_*ab*_, is defined as the probability that two alleles sampled at random from two individuals at the same autosomal locus are IBD. Let *k*_0*ab*_, *k*_1*ab*_, *k*_2*ab*_ denote the probability that individuals *a* and *b* share zero, one and two alleles IBD, respectively. The definition of kinship coefficient indicates that *ϕ*_*ab*_ can be expressed as a function of those IBD-sharing probabilities, to be more explicit, *ϕ*_*ab*_ = *k*_1*ab*_/4 + *k*_2*ab*_/2. Table 5 lists values of kinship coefficients, their corresponding IBD-sharing probabilities and the inference criteria of *ϕ*_*ab*_ derived using powers of 2 [18] for various relative pairs under the assumption of no inbreeding.

**Table 5.** Kinship coefficients for different relative pairs

| Relationship | $\phi_{ab}$ | $(k_{0ab}, k_{1ab}, k_{2ab})$ | Inference criteria |
| --- | --- | --- | --- |
| Monozygotic twins | 0.5 | (0, 0, 1) | $> \frac{1}{2^{3/2}}$ |
| Parent-offspring | 0.25 | (0, 1, 0) | $(\frac{1}{2^{5/2}}, \frac{1}{2^{3/2}})$ |
| Full sibs | 0.25 | (0.25, 0.5, 0.25) | $(\frac{1}{2^{5/2}}, \frac{1}{2^{3/2}})$ |
| Half sibs | 0.125 | (0.5, 0.5, 0) | $(\frac{1}{2^{7/2}}, \frac{1}{2^{5/2}})$ |
| Uncle-niece | 0.125 | (0.5, 0.5, 0) | $(\frac{1}{2^{7/2}}, \frac{1}{2^{5/2}})$ |
| First cousin | 0.0625 | (0.75, 0.25, 0) | $(\frac{1}{2^{9/2}}, \frac{1}{2^{7/2}})$ |
| Unrelated | 0 | (1, 0, 0) | $< \frac{1}{2^{9/2}}$ |

Suppose we have genotype data of *n* individuals, for each person we consider his/her genotypes at *m* SNP markers respectively. For 1 *≤ i ≤ n,* 1 *≤ j ≤ m*, let *X*_*ij*_ be the number of reference alleles (with label *A*) for individual *i* at SNP marker *j*. Thus *X*_*ij*_ takes values 0, 1, or 2 according to whether individual *i* has, respectively, 0,1, or 2 copies of allele *A* at marker *j*.

To simplify the illustration, we denote *μ*_*j*_ and 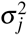 as the expectation and variance of *X*_*ij*_, respectively. In other words, 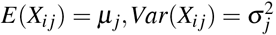. We assume the population variance for each marker is already known throughout our derivation. In practice, we can use sample variance, an unbiased estimator of population variance, as a substitute. Now we consider a pair of individuals *i* and *i*^′^. We use 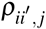 to denote the correlation coefficient between *X*_*ij*_ and 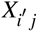. Besides, we let 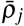 be the average of 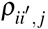 among all the individual pairs, i.e.

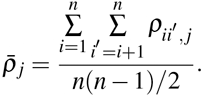

If we further assume all individuals are sampled from a homogeneous population, we can derive the following relationship among those correlations:

### Property 1.

Assume all individuals are sampled from a homogeneous population, then for 1 *≤ i, i^′^ ≤ n,* 1 *≤ j ≤ m*, we have

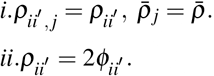

This property has also been mentioned in other articles, for example, see [11]. A proof of this property is given in S1 Appendix. Now we summarize the conclusions of this property as follows:

Result i. implies that the correlation between *X*_*ij*_ and 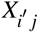 is irrelevant to which SNP we choose and depends only on the pair of individuals we select. Result ii. provides the quantitative relation between the kinship coefficient and the correlation of genotypes, which indicates that the estimation of kinship coefficient 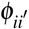 is equivalent to estimating the correlation coefficient of genotypes between individual *i* and *i*^′^ 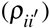.

Estimating kinship coefficient by calculating the average sample pairwise correlation among all genetic variants has been taken by many methods. Following this principle, a natural estimator of 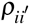 is

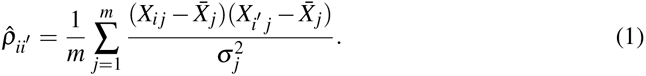

where 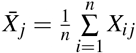 is the average counts of reference alleles (with label *A*) at SNP *j* in the whole population. We call 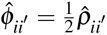 the scGRM estimator.

However, as we are going to demonstrate, 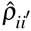 is actually a biased estimator of 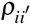. To illustrate this, we need the following property:

### Property 2.

For 1 *≤ i, i^′^ ≤ n,* 1 *≤ j ≤ m*, the estimated correlation coefficient between *X*_*ij*_ and 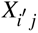 has a systematic bias from 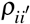. More specifically, we have

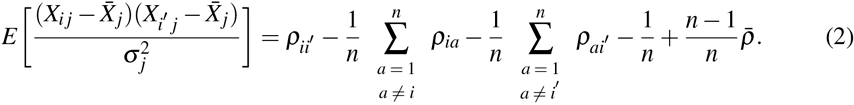

The proof is given in S1 Appendix.

Equation (2) also reveals that the expected value of 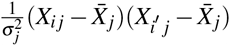 is not related to which SNP we select. Now we consider the expectation of estimator (1), it comes to the conclusion that

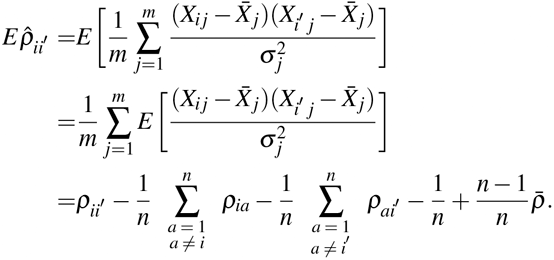

If 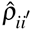 is an unbiased estimator of 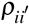, then we should have 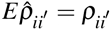. However, the result we derive is obviously contradictory to it. The existence of bias means a systematic error when we estimate kinship coefficient via the scGRM method mentioned above. To make this fact clearer, we sum the expectation of 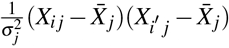 up over all the individual pairs in the population, which leads to the following property:

### Property 3.

For every SNP marker *j*, where 1 *≤ j ≤ m*, we have

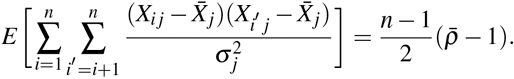

The proof is given in S1 Appendix.

Recall that 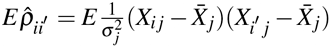, thus Property 3 also suggests

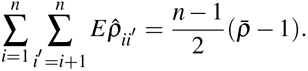

From Property 1 we know 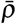 is the theoretical mean value of correlations between pair-wise individuals, therefore it must take the value between 0 and 1. This fact together with Property 3 reveals that the mean value of estimator 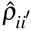 is negative on average, which explains the empirical observation that a substantial proportion of estimated kinship coefficients are negative.

This bias problem makes 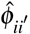 less desirable as an estimator of kinship between individuals *i* and *i*^′^. We can design an improved kinship estimation method which can eliminate the bias for each pair of individuals based on the scGRM estimator 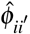. The improved estimation method, UKin, which stands for the unbiased kinship estimator, solves the bias problem without adding much computational complexity. To understand how this method guarantees the unbiasedness, we need the following property:

### Property 4.

For every SNP marker *j,* 1 *≤ j ≤ m*, and every pair of individuals *i* and *i*^′^, 1 *≤ i*, *i^′^ ≤ n*, we have

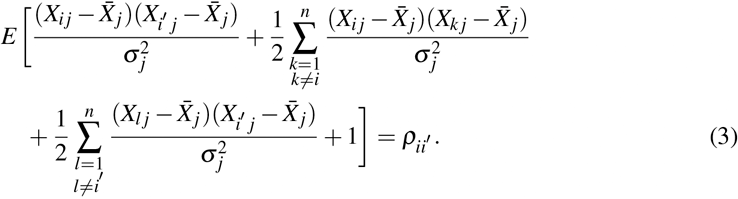

The proof is given in S1 Appendix.

For ease of presentation, we set

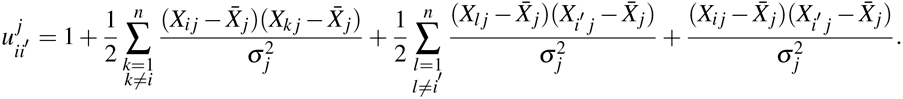

Using (3), we also conclude that the expectation of 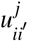 does not depend on which SNP we select. Based on this fact, a reasonable estimator of 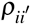 is

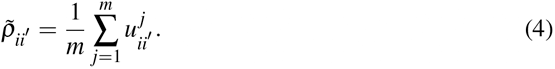

As Property 4 shows 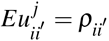 holds for every 1 *≤ j ≤ m*, the expectation of 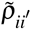 is still 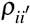. In other words, 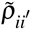 is an unbiased estimator of 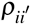, thus 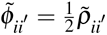 is an unbiased kinship estimator. Besides, as we can observe from the expression of 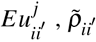 is the sum of a group of scGRM estimators 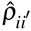 and a few correction terms, which means the UKin estimator relies on the same information we need for calculating the scGRM estimator 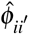. Thus the implementation of the UKin method doesn’t require extra data.

It is worth noting that there exists some relationship between the scGRM and UKin estimator. Substituting the expression of 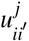 into (4), we get

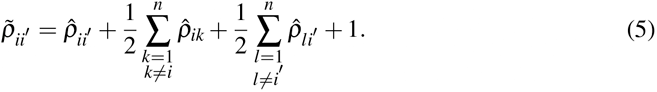

Equation (5) indicates that the UKin estimator 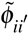 is a linear combination of some scGRM estimators 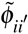 and constants. Thus 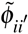 and 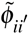 are based on the same genetic information. Besides, this conclusion also shows that the UKin method won’t bring a significant increase in computational complexity than the scGRM method.

Throughout our above analysis, we make assumptions of no inbreeding, LE and population homogeneity. In the Discussion we have analyzed these assumptions in detail.

## Supporting information

**S1 Appendix. Mathematical derivations of the properties in the Materials and Methods section.**

## Acknowledgments

This work was supported in part by the NIH grant GM134005 and NSF grants DMS 1713120 and 1902903.

## S1 Appendix

## Proof of Property 1.

For the *j*-th single nucleotide polymorphism (SNP) (1 *≤ j ≤ m*), let *f*_*j*_ be the frequency of the reference allele (with label *A*) at that SNP. Consider a pair of individuals *i* and *i*^′^ whose kinship coefficient is denoted by 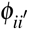, we derive the covariance of *X*_*ij*_ and 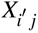 from two different aspects. Recall that we denote 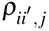 to be the correlation between *X*_*ij*_ and 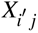, thus we have

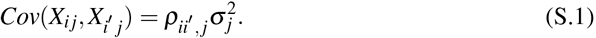

On the other hand, *X*_*ij*_ can be treated as the sum of two independent Bernoulli random variables. That is, *X*_*ij*_ = *B*_*ij*(1)_ + *B*_*ij*(2)_. For *k* = 1, 2,

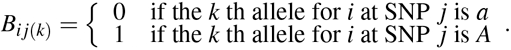

With this expression of *X*_*ij*_, we have

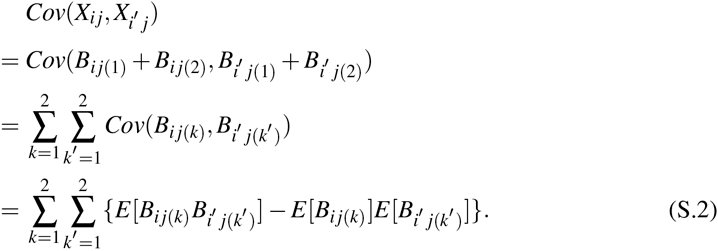

As we denote *f*_*j*_ to be the probability that a random allele chosen from the *j*-th SNP is *A*. Notice that 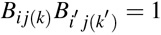 only when the two alleles selected from *i* and *i*^′^ at this marker are both with label *A*, under this circumstance, these two reference alleles are either identical by descent (IBD) or not. For simplicity, let *A*_*ij*_(*k*) represent the *k*-th alleles from individual *i* at SNP *j*, if we assume IBD genes have the same allelic types and non-IBD genes have independent allelic types, we obtain

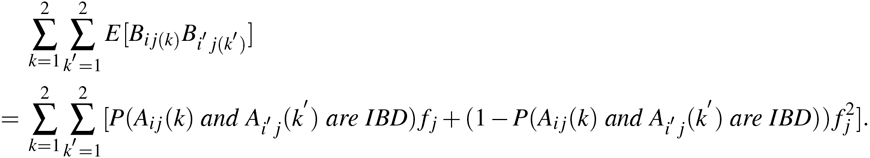

Consider the definitions of *f*_*j*_ and 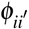, we obtain

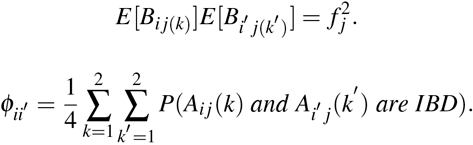

Substituting them into (S.2), we get

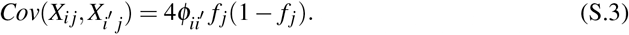

As *X*_*ij*_ is the sum of two *i.i.d.* Bernoulli random variables whose probability of success is *f*_*j*_, we can derive that 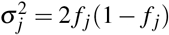. Together with (S.1) and (S.3), we have

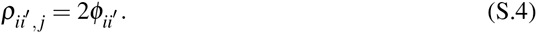

Equation (S.4) also reveals that the value of correlation 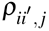 doesn’t depend on which SNP is selected, thus we get

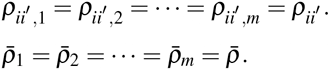

## Proof of Property 2.

To demonstrate this property, we need a few preparations:

i. Consider the result 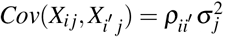, we have

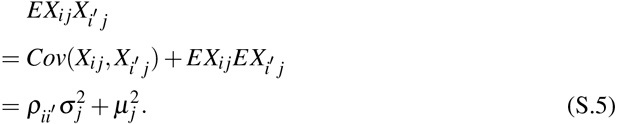 Directly applying this result yields

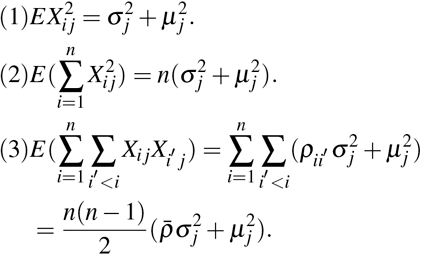
ii. Based on (1)–(3) stated above, we have

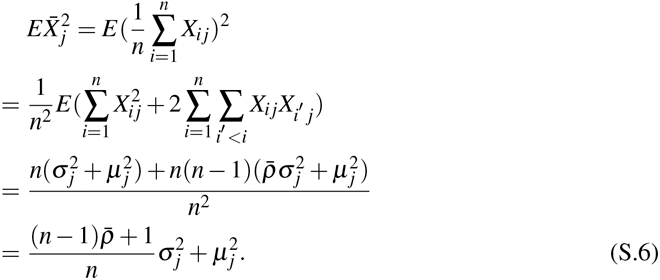

With these preparations, we now work on the demonstration of Property 2.

Directly expand the expression on the left side of (S.2), we have

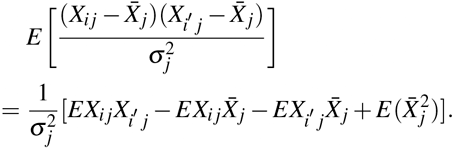

Substituting (S.5) and (S.6) into this expansion, we have

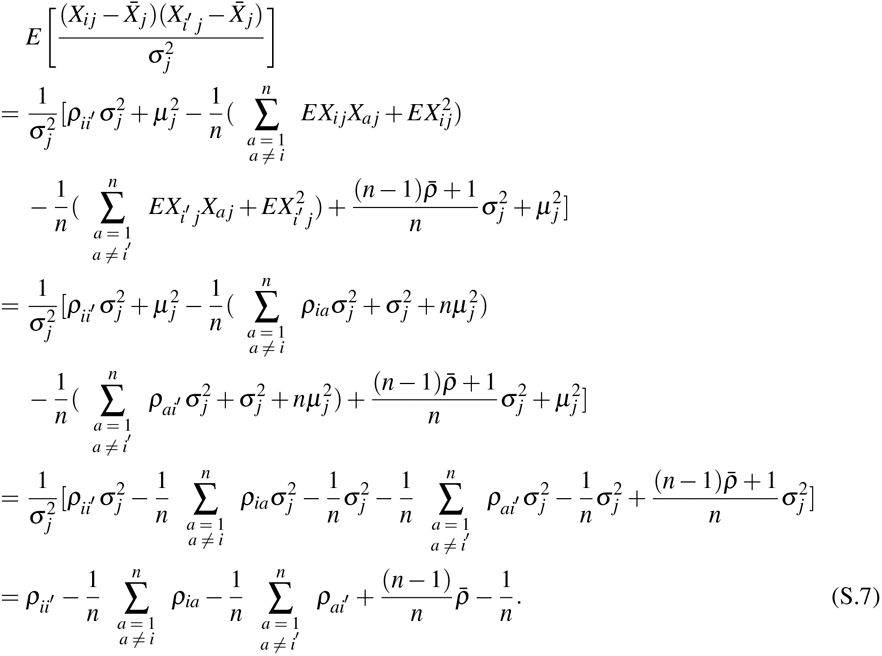

Thus we derive the conclusion in Property 2.

## Proof of Property 3.

For ease of calculation, we make a complement to the value range of index *i*^′^:

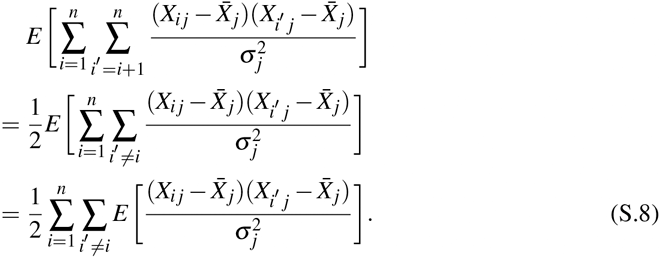

Equation(S.8) together with the conclusion (S.7) in the proof of Property 2 yields

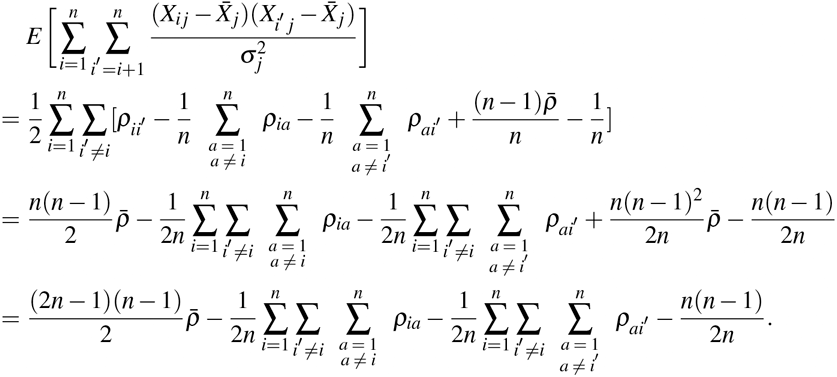

We observe that 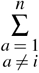 is irrelevant to *i*^′^, therefore

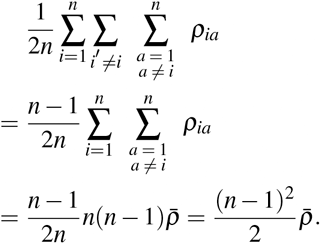

Besides, if we change the sequence of summation, we have

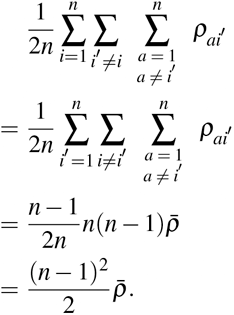

Substituting them into the expansion, we get

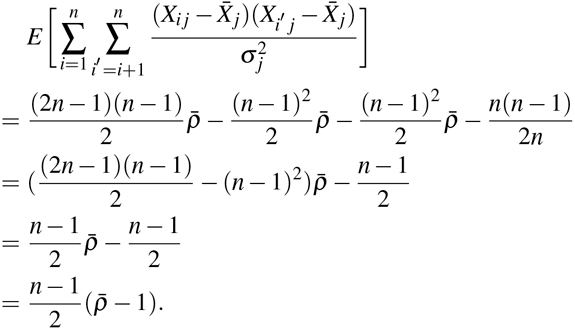

Thus we finish the proof of Property 3.

## Proof of Property 4.

At the start, we focus on a part of the expression on the left side of (3):

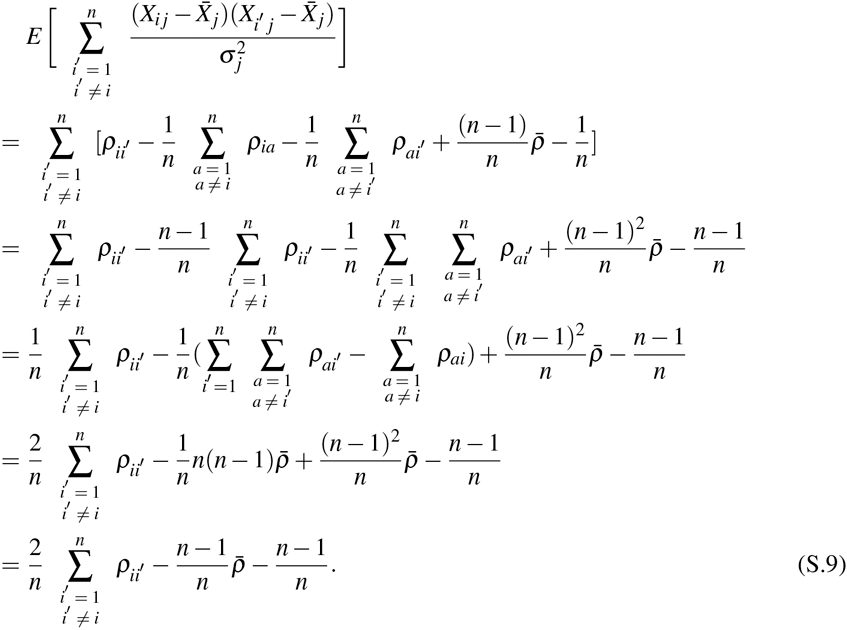

Substituting (S.7), together with (S.9), into the whole expansion, we get

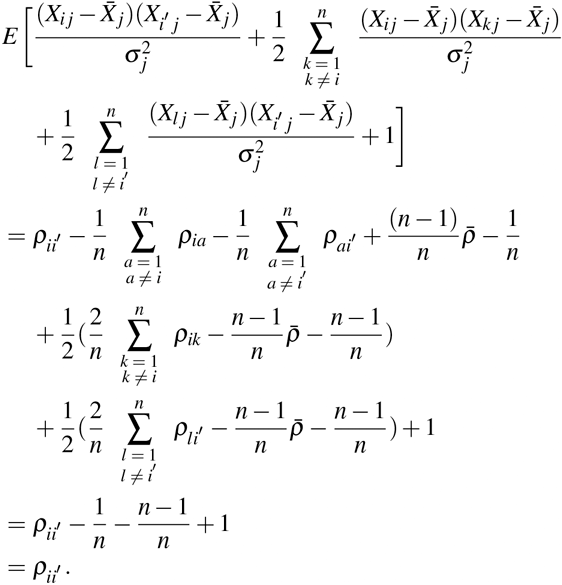

Here we have proved the conclusion in Property 4.

